# Exploring the effect of microdosing psychedelics on creativity in an open-label natural setting

**DOI:** 10.1101/384412

**Authors:** Luisa Prochazkova, Dominique P. Lippelt, Lorenza S. Colzato, Martin Kuchar, Zsuzsika Sjoerds, Bernhard Hommel

**Author notes:** authors contributed equally (shared first-authorship) (Luisa Prochazkova, Dominique P. Lippelt). **Corresponding author** L.Prochazkova, Leiden University, Faculty of Social and Behavioural Sciences, Cognitive Psychology Unit, Wassenaarseweg 52, 2333 AK Leiden, The Netherlands.

## Abstract

**Introduction:** Recently popular sub-perceptual doses of psychedelic substances such as truffles, referred to as microdosing, allegedly have multiple beneficial effects including creativity and problem solving performance, potentially through targeting serotonergic 5-HT_2A_ receptors and promoting cognitive flexibility, crucial to creative thinking. Nevertheless, enhancing effects of microdosing remain anecdotal, and in the absence of quantitative research on microdosing psychedelics it is impossible to draw definitive conclusions on that matter. Here, our main aim was to quantitatively explore the cognitive-enhancing potential of microdosing psychedelics in healthy adults.

**Methods:** During a microdosing event organized by the Dutch Psychedelic Society, we examined the effects of psychedelic truffles (which were later analyzed to quantify active psychedelic alkaloids) on two creativity-related problem-solving tasks: the Picture Concept Task assessing convergent thinking, and the Alternative Uses Task assessing divergent thinking. A short version of the Ravens Progressive Matrices task assessed potential changes in fluid intelligence. We tested once before taking a microdose and once while the effects were manifested.

**Results:** We found that both convergent and divergent thinking performance was improved after a non-blinded microdose, whereas fluid intelligence was unaffected.

**Conclusion:** While this study provides quantitative support for the cognitive enhancing properties of microdosing psychedelics, future research has to confirm these preliminary findings in more rigorous placebo-controlled study designs. Based on these preliminary results we speculate that psychedelics might affect cognitive metacontrol policies by optimizing the balance between cognitive persistence and flexibility. We hope this study will motivate future microdosing studies with more controlled designs to test this hypothesis.

## Introduction

Major news outlets throughout the world are reporting on the growing number of professionals using small doses of psychedelics (e.g. magic mushrooms, truffles, or peyote) to boost their productivity and creativity at work. A prominent example is the use of small doses of LSD by employees in Silicon Valley, as a ‘productivity hack’ (Glatter 2015). This emerging phenomenon is referred to as microdosing, with dosages around one tenth of recreational doses. Yet, despite the low dosages microdosing is still thought to provide a potential boost in cognition according to anecdotal reports (Cooke 2017; Gregorie 2016; Leonard 2015; Sahakian 2017; Senior 2017; Solon 2016). Moderate to large doses of psychedelics induce changes in perception, mood and overall consciousness, often described as qualitatively similar to deep meditative or transcendental states (Barrett and Griffiths 2017; Barrett, Johnson and Griffiths 2017). If, similar yet milder effects apply to microdosing, this would render microdosing a potentially interesting cognitive enhancer in healthy individuals or even the basis of a treatment strategy to tackle various disorders including depression.

Throughout the 1960’s psychedelics were extensively used at recreational doses in experimental research, clinical settings, and in creative and scientific vocations (Sessa 2008), but were made illegal in most countries worldwide as a reaction to the rising counterculture of the 1960’s and failure to establish the clear efficacy of LSD treatment (Oram, 2014). Now, after many decades of disregard, psychedelics have started to reappear as a genuine and promising area of research within experimental and clinical psychology, as well as psychiatry. Moreover, certain psychedelics, such as truffles, have regained a legal status in The Netherlands, offering researchers a particularly interesting opportunity to study its effects in a quantitative manner. This is highly desirable, as previous reports have remained anecdotal (Oberhaus 2017) and qualitative at best, often focusing on experiences of elevated feelings of determination, alertness, and energy, improved pattern recognition, as well as strong reductions of depressive feelings (Fadiman and Krob 2017). Qualitative studies based on self-reports are known to suffer from validity problems due to participants’ inaccurate memories, differences in vocabulary and verbal skills, and unintentional or willful distortions of subjective experiences (Schwarz 1999).

Nonetheless, existing research with moderate doses of psilocybin shows that psilocybin is a potent neuro-pharmacological agent with a strong modulatory effect on brain processes (Vollenweider and Kometer 2010). Furthermore, a double-blind placebo-controlled study by Hasler et al. (2004) showed that even very low doses of psilocybin (45 µg/kg body weight) were rated clearly psychoactive by most of the volunteers, which indicates that psychedelic effects do not need high doses to be recognized.

Classical psychedelics such as psilocybin (O-phosphoryl-4-hydroxy-N, N-dimethyltryptamine), the active compound in psychedelic truffles, exert their primary effects by directly binding to serotonin 2A receptors (5-HT_2A_; Tylš et al. 2014). Interestingly, 5-HT_2A_ agonism has been reported to be associated with enhanced cognitive flexibility (Boulougouris and Robbins 2010), improved associative learning (Harvay 2003) and hippocampal neurogenesis (Catlow 2016) in animals. Additionally, psychedelics have been shown to increase subjective sense of wellbeing, optimism, and openness in humans (Griffiths et al. 2006, 2008; MacLean et al. 2011). Moreover, multiple clinical trials using moderate to large doses of psychedelics have indicated that psychedelics have anxiolytic, antidepressant (dos Santos et al. 2016; Carhart-Harris et al. 2016; Grob et al. 2011; Griffiths et al. 2016), anti-compulsive (Moreno et al. 2006), and anti-addictive properties (Bogenschutz et al. 2015, Johnson et al. 2014, dos Santos et al. 2016). Consequently, the effects of psychedelic substances can be argued to target the serotonergic system, and hence be beneficial in situations where there is need for mental flexibility, or where one needs to break through rigid patterns of thought. In case that future research confirms positive effects of microdosing on brain and cognition, microdosing could become an attractive alternative due to its sub-perceptual nature possibly sparing individuals from the perceptual distortions often reported with moderate or high doses.

Through the alleged benefits in mental flexibility, a promising behavioral target of psychedelics lies in the area of creativity. Creativity is a multilayered phenomenon, commonly defined as the ability to generate ideas, solutions, or products that are both novel and appropriate (e.g., Amabile 1996; Sternberg and Lubart 1999). Creativity is not a unitary function but consists of a number of subcomponents (Wallas 1926) that provide different, to some degree opposing cognitive challenges. It is crucial to distinguish between convergent thinking, which requires identification of a single solution to a well-defined problem (Mednick 1962), and divergent thinking, which requires the collection of many possible solutions to a loosely defined problem (Guilford 1967). An example of convergent thinking task would be to find the one concept that can be meaningfully combined with three other concepts such as “…man”, “…market”, and “…bowl” (such as “super”), while an example of divergent thinking task would be to list all possible ways in which a brick could be used (for throwing, as a weight, as a weapon, etc.). It has been argued that convergent thinking draws more on the ability to focus exclusively on a given problem (persistence), while divergent thinking draws more on cognitive flexibility (Lippelt, Hommel, and Colzato 2014). However, it is important to point out that all available creativity tasks require the integration of both of these abilities to some degree. Of further importance to our present study is the fact that creative thinking is not a hardwired virtue. Several behavioral studies have shown that the processes underlying creative thinking can be systematically enhanced and impaired by both behavioral interventions, such as meditation, as well as, psychopharmacological agents as for instance cannabis, tyrosine and Adderall (e.g., Baas, Nevicka, and ten Velden 2014; Colzato de Haan and Hommel 2015; Farah et al. 2009; Kowal et al. 2015; Ritter and Mostart 2016; Schafer et al. 2012; Zabelina and Robinson 2010; Davis 2009).

Moreover, a recent study conducted by Kuypers and colleagues (2016) investigated the effect of recreational doses of the psychedelic brew Ayahuasca on creativity during two spiritual retreats. They found that divergent thinking performance improved under the influence of Ayahuasca compared to baseline, while convergent thinking performance decreased in comparison to baseline. Although this study may seem to provide a useful starting point, conclusions are hampered by several disadvantages of this drug and the study design. First, dimethyltryptamine (DMT), the active psychedelic compound in Ayahuasca, needs to be combined with monoamine oxidase inhibitors (MAOIs) for its effect to take place. MAOIs are known to have anti-depressant effects on their own, so they represent a possible confound in all Ayahuasca studies (Quitkin 1979). Additionally, this implies that the qualitative experience induced by Ayahuasca and the underlying mechanisms of action differ substantially from those related to commonly used psychedelics, such as LSD or psilocybin (Callaway 1998; Riba et al. 2006). Indeed, the Ayahuasca brews used in the study of Kuypers and colleagues’ (2016) induced strong psychedelic experiences, the effects of which are unlikely to be comparable to the effects obtained from microdosing a psychedelic substance. High doses of psychedelics frequently result in disorienting effects in the user, which makes reliable assessment of psychometric task performance difficult during the peak effects of the psychedelic experience (Hollister 1968). Taken altogether, microdosing of psychedelic truffles and related psychoactive substances may thus be more suitable to assess the enhancing effects of psychedelics on human performance.

The current study is the first to experimentally investigate the cognitive enhancing effects of microdosing on human cognition in a natural setting. We were offered the unique opportunity to quantitatively study the effects of microdosing truffles during microdosing events of the Psychedelic Society of The Netherlands (PSN), a non-laboratory environment. Natural settings like such an event have a number of potential benefits as they are more comparable to situations of real life use than studies in a laboratory environment. The truffle samples, which participants obtained during the microdosing event, were post-hoc analyzed, to determine the exact amount of active substance potentially leading to the found effects. The major aim of our study was to study the effects of psychedelic truffles on creative thinking. We assessed convergent and divergent thinking separately, by using the Picture Concept Task (PCT; Hurks et al. 2010; Wechsler 2003) and Guilford’s (1967) Alternate Uses Task (AUT), respectively. Given that convergent thinking is correlated with fluid intelligence (e.g., Akbari Chermahini, Hickendorff and Hommel, 2012), we also employed a short 12-item version of Raven’s Progressive Matrices Task (Bilker et al. 2012), a standard intelligence test, once before and once during the acute effects of a microdose of truffles. Given the effects of large doses of psychedelics on positive mood, trait openness, and assumed cognitive flexibility reflected by psychosis-like symptoms (Carhart-Hariss et al. 2016), we expected improvements on the AUT after microdosing, as mood and flexibility are two factors that are known to boost divergent thinking (Baas, de Dreu and Nijstad 2008; de Dreu, Baas and Nijstad 2008; Zabelina and Robinson 2010). Due to a lack of relevant previous studies and the subjectivity of self-reports, the effect on convergent thinking was difficult to predict. On the one hand, previous dissociations of convergent and divergent thinking might suggest that convergent thinking is impaired by microdosing—an outcome that could imply that microdosing shifts cognitive control states from persistence to flexibility (Hommel 2015). However, microdosing may also improve both convergent and divergent thinking, rather suggesting that microdosing improves the interplay between persistence and flexibility. We did not have specific expectations regarding intelligence, but were interested to see whether possible effects on convergent thinking might generalize to performance on the intelligence task, or remained more specific.

## Methods

### Procedure

The experiment was conducted during a microdosing event organized by the Psychedelic Society of the Netherlands (PSN), who provided us with the opportunity to ask participants to take part in the experiment by means of an on-stage presentation. Interested participants were presented with envelopes containing the informed consents. We asked participants to read the provided information carefully, to sign it if they gave permission for their participation and use of their anonymized, coded data, and to subsequently return it back in the envelope. The envelope also included the experimental tasks for the first session as a booklet. However, we stressed participants to not open the booklet until one of the experimenters asked them to turn the first page to prevent premature exposure to the tasks. The experimenters also kept a close eye on the attendees to further ensure participant compliance. Next, all participants were carefully guided through the experimental tasks. Each task was explained in detail by one of the experimenters with the aid of example items, before participants were allowed to perform the task themselves. This was repeated for each of the three tasks. The protocol was approved by the local ethics committee (Leiden University, Institute of Psychology). The experiment consisted of a baseline session before participants had consumed any psychedelics and a second session carried out while participants were under the influence of a microdose of psychedelic truffles. The tasks were conducted in a group setting free from outside distraction during both sessions. Participants’ responses were assessed in paper-and-pencil version. The test battery consisted of the Picture Concept Task (PCT; Wechsler 2003; Hurks et al. 2010) to assess convergent thinking, the Alternate Uses Task (AUT; Guilford 1967) to assess divergent thinking, and a validated short 12-item version of the Raven’s Progressive Matrices Task (RPM; Bilker et al. 2012) to test fluid intelligence. While AUT and RPM were presented fully in paper version, PCT stimuli were presented by PowerPoint to insure precise timing of stimuli presentation (further details about the contents of the psychometric tasks are provided under the “Materials” section). Pre- and post-microdosing performance was assessed by administering two different versions of each task, to reduce potential learning effects. Task versions were counterbalanced across sessions and participants.

After finishing the experimental tasks in the first session, participants consumed a microdose of premeasured psychedelic truffles made available by the PSN. Participants of the workshop agreed to take psychedelic truffles upon their own risk. The PSN did not adhere to any strict guidelines regarding the dosage given to attendees. Nonetheless, they did take the attendees self-reported approximate weight into account in their recommended dosage, by means of subjective evaluations regarding low, average and high body weight criteria. Attendees judged to have a low body weight were recommended to take 0.22 grams of truffles, those with an average body weight 0.33 grams, and those with a high body weight were recommended 0.44 grams of dried truffles. Moreover, it should be noted that attendees did not have to adhere to the PSN member’s recommendation and were free to choose a dose. However, with reference to the existing microdosing guidelines (Fadiman, 2011) the doses provided to participants appeared to be in a meaningful range of a microdose. Following the publication of The Psychedelic Explorer’s Guide (Fadiman, 2011) a microdose should lie around 1/10th to 1/16th of a regular dose. Considering that a recreational dose of truffles is about 10 grams of fresh truffles, a microdose would equal 1 gram of fresh truffles. As fresh truffles consist of 2/3rds of water, this results in a weight of 0.33g of dried truffles. Participants consumed on average 0.37 of dried truffles which is an appropriate amount given the calculation. Additionally, data on participants’ height, body weight, and ingested dose of truffles were independently collected by the researchers in order to examine potential dose dependent effects in the analysis.

Approximately 1.5 hours after attendees had consumed the truffles, participants were asked to take part in the second session of the experiment. The 1.5-hour time interval was chosen as the effects of truffles are reported to peak around 30-90 minutes followed by a few hours long plateau of effects before rapidly subsiding back to baseline (Erowid 2017). By choosing this time interval we could be certain that all participants were tested while the effects of the truffles were still plateauing during the second session. The procedure during the first session was repeated at the second session. Lastly, participants filled in a questionnaire on medical health, psychedelic and general drug use, and general personal information (e.g. gender, age, first language). After testing, we thanked participants for their participation and debriefed them about the purpose of our experiment.

### Sample

Out of all 80 attendees at the microdosing event, 38 volunteered in our experiment. Participants indicated that they were healthy, spoke the required languages for performing the experimental tasks, and whether they had prior experience with the use of psychedelic substances. All 38 participants completed the RPM during both sessions. Regarding the PCT, eleven participants had to be excluded from further analysis, either due to incorrect interpretation of the task instructions or because they did not complete the task either for the first or second session. For the AUT, two participants had to be excluded as the experimenters were unable to read the individuals’ writing, two due to missing data on the second session, and one due to misinterpretation of the task instructions. Importantly, the individuals that were excluded for the AUT and PCT analysis did not overlap. This suggests that exclusion was random and did not depend on a shared trait or state among those who were excluded from the two analyses. Nonetheless, in order to preserve power we decided to analyze the data separately for each task. This yielded a sample of 38 participants for the analyses of the RPM data, 27 subjects for the PCT analyses and 33 for the analyses of the AUT data. Table 1 provides an overview of additional descriptive information regarding the sample.

**Table 1:** Descriptive statistics for the samples used in the three separate analyses. Numbers indicate Mean (SD), unless otherwise specified (Gender & Prior Experience)

| Sample | Age | Gender<br>(M/F) | BMI | Weight<br>(kg) | Prior<br>Experience<br>(Y/N) | Ingested<br>Dosage (g) |
| --- | --- | --- | --- | --- | --- | --- |
| **RPM** | 31.1 (11.49) | 23 / 15 | 22.5 (4.74) | 68.4 (12.81) | 36 / 2 | 0.37 (0.159) |
| **AUT** | 30.0 (10.89) | 19 / 14 | 22.4 (4.92) | 67.5 (13.01) | 31 / 2 | 0.35 (0.142) |
| **PCT** | 31.5 (12.49) | 18 / 9 | 22.9 (5.42) | 69.3 (13.36) | 25 / 2 | 0.41 (0.158) |

### Truffles analysis

Dried truffle samples of what participants used during the microdosing event (corresponding to 0.22, 0.33, or 0.44 grams dry weight) were post-hoc analyzed at the University of Chemistry and Technology Prague (UCT; Laboratory of Biologically Active Substances and Forensic Analysis) to determine the exact amount of active substances potentially leading to the found effects. Analytical standards of psilocin, psilocybin, norbaeocystin, and baeocystin were synthesized in-house (purity ≥95%) at UCT. The evaluation of the developed analytical method (Hajkova et al. 2016) encompassed the determination of Limit of Quantitation (LOQ) and the determination of the applicable concentration range for each compound studied. LOQ was determined as 10 times the ratio of signal to noise. Concentration was determined for each of the truffle samples separately and each sample was measured twice. The level of the alkaloids was almost identical in all three samples, however, and the differences were lower than the estimated measurement errors when the results from the different samples is averaged. As such, we report the concentrations for the four alkaloids collapsed across the three samples in the results. For the alkaloid concentrations for the three samples separately and further details regarding the methods underlying these analyses please see Online Resource 1

Despite psilocybin usually being the most abundant alkaloid in psychedelic truffles psilocin is commonly considered to be responsible for the psychedelic truffles’ characterizing psychoactive effects (Gartz et al.,1994; Tyls et al., 2014). Psilocybin is quickly metabolized into psilocin in the human body through dephosphorylization, thereby limiting psilocybin’s direct contribution to the psychoactive effects in humans. Furthermore, we also report the concentrations of baeocystin and norbaeocystin as it has recently been suggested that they could quickly metabolize into psilocin upon consumption through similar processes as are at play in the metabolism of psilocybin (Tyls et al., 2014).

### Instruments

#### Picture Concept Task

The PCT (Wechsler 2003; Hurks et al. 2010) is a visual creativity task that involves finding a common association between several images. Each trial consists of a matrix of between 2×3 to 3×4 pictures (see Fig 1 for an example). The correct solution is a common association between one picture from each row. Thus, a response indicating an association between two pictures from the same row would be incorrect. Because the task presumes there is only one correct solution to each item, the task has been previously used to measure convergent thinking (Hurks et al. 2010). Hence, to complete the task one should converge on the correct solution, while inhibiting inappropriate or less obvious associations and previously attempted yet incorrect solutions.

**Figure 1:**
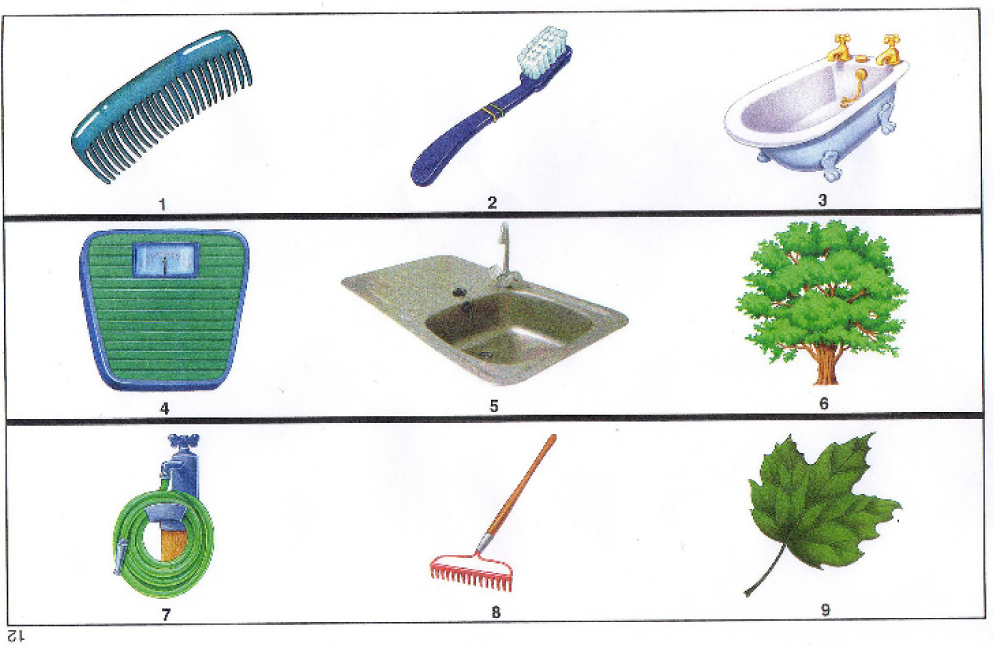
Example trial from the Picture Concept Task. The subject has to identify the common association between 1 item from each row

Participants had 30 seconds per item to find the solution. Because precise time limit is essential in this task, it was impractical to present the task on paper. Instead, we used a PowerPoint presentation in which the slides (i.e. items) transitioned every 30 seconds. Participants were instructed to mark and name the common association between the pictures on the PCT response sheet in the booklet (see Fig 2 for a corresponding answer sheet belonging to the item in Fig 1). The PCT was scored by summing the number of correct responses.

**Figure 2:**
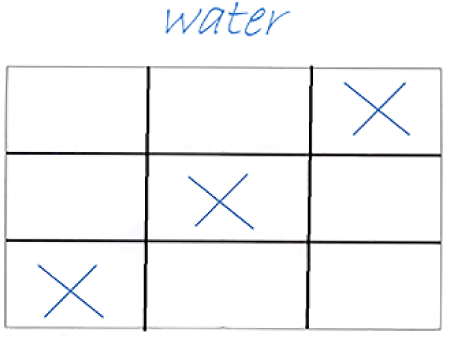
Illustration of a filled in answer sheet for the items from Fig. 1

#### Alternate Uses Task

The AUT is commonly used in research on creativity to measure divergent thinking performance (Guilford 1967). During the AUT subjects are presented with a common household object and asked to think of as many possible uses for the object as they can within a limited amount of time. Within each session of our experiment participants were either presented with the word Pen or the word Towel and given 5 minutes (per session) to write down as many possible uses for the object. As common, the AUT was rated according to four different variables:

- Fluency: the total number of responses
- Flexibility: the number of different categories of responses
- Elaboration: how much the person elaborates on their response. Each ‘elaboration’ receives one point. For instance, the response “using a brick to prevent a door from slamming shut (1), when it is windy (2)” earns 2 elaboration points.
- Originality: the uniqueness of a response. Originality is calculated by dividing the total number of responses by *all* subjects, once by 5% and once by 1%. Responses that have also been mentioned by 1% or less of the other participants receive 2 points for originality, while responses that have been mentioned by 1-5% of the participants receive 1 point for originality. All other responses receive no originality points.

Out of these four, flexibility is the most reliable and theoretically most transparent index of divergent thinking (Akbari Chermahini and Hommel 2010), while fluency neglects the quality of responses and the originality score is highly sample-dependent. As the scoring of AUT variables can be highly subjective we used the mean score for each of the four measures obtained from by two independent raters to increase the reliability of the dependent measures. Reliability scores (Cohen’s κ) were very high for fluency (Session 1: κ = .970; Session 2: κ = .935), fair for flexibility (Session 1: κ = .342; Session 2: κ = .252), fair to moderate for elaboration (Session 1: κ = .231; Session 2: κ = .547) and fair for originality (Session 1: κ = .254; Session 2: κ = .318).

#### Raven’s Progressive Matrices

The RPM was developed by Raven (1938) in order to measure fluid intelligence. We used a shortened 12- item version of the RPM to reduce testing time and participant burden. The 12-item version was developed and validated by Bilker and colleagues (2015) showing high correlations with the full RPM (*r*= .80 for version A, and *r* = .77 for version B).

In our experiment, the RPM consisted of a series of 2×2 or 3×3 matrices of pictures in which the lower right picture was always missing. Both horizontally and vertically, a pattern is present in the matrix of pictures allowing the participant to deduce what the missing picture should look like. Underneath each item 6 possible solutions were presented and participants marked the correct solution by circling it on paper. While there was no time limit per item, the task had a total time limit of 5 minutes. Throughout the RPM items increased in difficulty, but participants were allowed to skip an item in case they felt like they were stuck. However, once they had advanced to a next item (either by skipping or answering) they were no longer allowed to go back to correct earlier responses. The RPM was also scored by summing the number of correct responses.

#### Analyses

First, separate analyses were run to test for possible interactions between time-point (pre vs. post truffle ingestion) and participants’ body weight, ingested dose, and prior experience with psychedelic substances on the dependent measures. As we did not find any significant interactions we dropped these factors from any subsequent analyses. To assess changes in fluid intelligence, we performed a paired samples t-tests comparing RPM scores at baseline versus RPM scores post ingestion of the truffles in the entire sample. Next, we performed a paired samples t-test to compare convergent thinking performance before and after ingestion for the 27 participants for whom we obtained valid data on the PCT for both sessions. To assess microdosing-induced changes in divergent thinking performance we analyzed the variables obtained from the AUT. Although of main interest were the flexibility and fluency scores, for completeness we entered all four variables (i.e. fluency, flexibility, elaboration and originality) as separate dependent measures in one multivariate Repeated Measures ANOVA with time (pre-versus post-ingestion) as the within subject factor. Significant effects in the multivariate ANOVA were followed-up by the appropriate univariate tests. The significance level for all analyses was set to α = .05.

## Results

### Psychedelic content in truffle sample

The Forensic Laboratory of Biologically Active Substances confirmed the presence of the active psychedelic alkaloids in the truffle samples. Quantitative analysis showed the presence of all four investigated alkaloids with predominant levels of psilocybin (see Table 2 for an overview of alkaloid concentrations). The next-most abundant alkaloids were psilocin followed by baeocystin. The concentrations of the four alkaloids were almost identical for all three samples and the differences were lower than the measurement errors obtained when the data is collapsed across samples.

**Table 2:** Results of observed analytes averaged over the three dosages

| Alkaloid | µg/g (ppm) | SD | Relative SD |
| --- | --- | --- | --- |
| Psilocybin | 1595 | 3.75 | 2% |
| Psilocin | 85 | 0.86 | 10% |
| Norbaeocystin | 8 | 0.06 | 7% |
| Baeocystin | 31 | 0.15 | 5% |

In this study, we compared task performance before the ingestion of the psychedelic truffles with performance approximately 1.5 hours after ingestion (i.e. while the effects of the psychedelic truffles were manifested). Table 3 displays descriptive statistics for both sessions of performance on all dependent measures.

**Table 3:** Descriptive statistics (Means [SE]) for the first and second session

| Measure | Session 1 | Session 2 |
| --- | --- | --- |
| PCT | 6.56 (1.601) | 7.59 (1.600) |
| RPM | 8.58 (2.238) | 8.97 (1.924) |
| Fluency | 14.68 (0.99) | 16.70 (1.19) |
| Flexibility | 11.20 (0.80) | 12.74 (0.93) |
| Elaboration | 2.18 (0.32) | 1.76 (0.28) |
| Originality | 12.36 (1.30) | 15.67 (1.45) |

### Interaction effects with weight, body-mass-index, ingested dosage and prior experience

We tested for possible interaction effects on all dependent measures using Repeated Measures ANOVAs. All *F*s were smaller than 1, except for the interaction with the ingested dosage on the PCT, *F*(1, 25) = 1.13, *p* = .299, and the interactions with prior experience on the PCT, *F*(1, 25) = 1.05, *p* = .316, and RPM, *F*(1, 36) = 3.69, *p* = .063. Nevertheless, these results show that none of the factors significantly interacted with the independent factor time-point and were thus dropped from the final analyses described below.

### Fluid Intelligence

Comparing fluid intelligence on pre- and post-microdosing using a paired-samples t-test, we found no difference between the two time-points with respect to the number of correct items on the RPM, *t*(37)=1.00, *p*=.324, Cohen’s *d*=.163.

### Convergent thinking

Performance on the PCT (number of correct responses) was significantly higher in the second than in the first session, *t*(26)=2.56, *p*=.017, Cohen’s *d*=.493, showing an improvement of convergent thinking.

### Divergent thinking

The four measures of the AUT were analyzed by means of a multivariate Repeated Measures ANOVA. The effect of time-point was significant, *F*(4, 29)=4.16, *p*=.009, partial η^2^=.365, due to better performance in the second than the first session. Additional univariate tests showed a significant increase in fluency, *F*(1, 32)=5.59, *p*=.024, partial η^2^=.149, flexibility, *F*(1, 32)=6.23, *p*=.018, partial η^2^=.163, and originality scores, *F*(1, 32)=12.03, *p*=.002, partial η^2^=.273, while the change in elaboration scores failed to reach significance, *F*(1, 32) = 2.97, *p* = .226, partial η^2^ = .046.

## Discussion

The aim of this study was to explore the effects of microdosing psychedelics on creative problem solving. We observed an increase in divergent idea generation on the AUT, as evidenced by a significant increase in fluency, flexibility, and originality scores, as well as an increase in convergent thinking on the PCT after intake of a microdose of magic truffles. Given that fluid intelligence did not change between the two measurement time points suggests a specific effect on creativity performance, but not on general cognition. These findings are in line with earlier studies finding positive effects of high doses of psychedelics on creative performance (Harman and Fadiman 1970; Harman et al. 1970; Kuypers et al. 2016; Zegans et al. 1967). In particular, the increase in originality scores on the AUT parallels the increase in originality scores after intake of Ayahuasca reported by Kuypers and colleagues (2016). Taken together, our results suggest that consuming a microdose of truffles allowed participants to create more out-of-the-box alternative solutions for a problem, thus providing preliminary support for the assumption that microdosing improves divergent thinking. Moreover, we also observed an improvement of convergent thinking, that is, increased performance on a task that requires the convergence on one single correct or best solution.

Before continuing to interpret our findings, it is important to consider the implications of the fact that we did not use a control group (for obvious ethical and practical reasons). Given the absence of a control group, we cannot rule out the possibility that changes from the first to the second time point of measurement are due to the impact of other factors than the microdosed truffles. Two of these factors come to mind. For one, it is possible that increased performance from the first to the second time point of measurement reflects learning. We consider that possibility not very likely, for three reasons. First, it does not seem to fit with the absence of an improvement for the intelligence measure, even though the Raven task shares many aspects with the PCT and the AUT. Second, studies on convergent thinking have not shown evidence of improved performance with multiple testing—at least if different test items were used. For instance, Colzato, Ozturk, and Hommel (2012) had participants perform the Remote Association Task, which can be considered a verbal version of the PCT, three times in different conditions and found neither condition effects nor, and this would be the learning-sensitive test, any interaction between condition and condition order. Third, a recent training study did not reveal any (positive) training effects on AUT performance (Stevenson, Kleibeuker, de Dreu, and Crone, 2014). No effect of 8 training sessions was observed for originality, a *negative* training effect was obtained for fluency, and a quadratic effect for flexibility. Both fluency and flexibility measures in fact *decreased* over the first 3 to 4 sessions, and only the flexibility measure eventually reached the original baseline in the 8^th^ session. Taken together, we see no empirical support for the possibility that our observations might reflect a learning effect.

For another, it is possible that increased performance from the first to the second time-point reflects an effect of expectation. Expectation effects are widely studied but not well understood (Schwarz, Pfister, and Büchel, 2016). Drug-related expectation effects commonly require previous experience with the psychological effects of the respective drug, and it is likely that expectations operate by having been associated with, and thus conditioned to stimuli and expectations preceding the actual effect (Schwarz et al., 2016). If so, the existence of expectation-based effects does not contradict the existence of real drug effects, and the former in fact rely on the previous experience of the latter. From that perspective, expectation-based effects and drug-induced effects are likely to have comparable impact on psychological functions, presumably even through the same physiological means. Accordingly, while we cannot definitely exclude that the effects we observed actually required the present intake of the drug, they even in the worst case are likely to rely on the previous intake and likely to reflect the effects of that previous intake.

Notwithstanding these caveats, the outcome pattern of the present study is consistent with the idea that microdosing psychedelic substances improves both divergent and convergent thinking. The fact that intelligence was not improved suggests that this effect was rather selective, but the possibility remains that the Raven was less sensitive to the intervention than the other measures were. It is tempting to interpret our observations on divergent thinking in the context of recent suggestions that behavior drawing on flexibility and novelty benefits from a reduction of cognitive top-down control (Cools and D’Esposito 2009; De Dreu et al., 2008; Dreisbach and Goschke 2004; Hommel 2015; Nijstad et al., 2010). According to this view, creativity tasks can be assumed to draw on two distinct, presumably opposing cognitive processes: flexibility is characterized by broadening the attentional scope, which enables individuals to generate many divergent ideas, while persistence is associated with a narrower attentional scope, thus allowing individuals to focus on one creative idea at a time (De Dreu et al., 2008; Hommel 2015). Some of the previous empirical dissociations of persistence and flexibility were related to dopaminergic functioning, such as in behavioral-genetics studies demonstrating that polymorphisms supporting efficient dopaminergic functioning in the frontal cortex promote persistence while polymorphisms supporting striatal dopaminergic functioning promote flexibility (e.g., Reuter, Roth, Holve and Hennig, 2006; Zabelina, Colzato, Beeman and Hommel, 2016; for an overview, see Hommel and Colzato, 2017). This strengthens the view that frontal and striatal dopaminergic pathways are involved in persistence and flexibility. If we assume that convergent thinking relies more on frontal persistence while divergent thinking relies more on striatal flexibility, our outcomes raise the question how an intervention can manage to improve both convergent and divergent thinking.

Classical hallucinogens, including psilocybin, belong to a group of tryptamines that are thought to exert their primary psychedelic effects through activity at the serotonergic 5-HT_2A_ receptor (Vollenweider and Kometer 2010). Of particular interest in this regard are findings from animal studies showing that 5- HT_2A_ agonist activity (Halberstadt 2015) correlates with an increase in associative learning (Harvey et al. 2003) and improvements in the ability to adapt behavior more flexibly (Boulougouris et al. 2008). Moreover, studies in humans have shown that the administration of psychedelics is associated with an increase in the personality trait “Openness” (MacLean et al. 2011) and that psychedelics can induce a reduction in symptoms associated with rigid behavior and thought patterns observed in obsessive-compulsive disorder (Moreno et al. 2006) and depression (Carhart-Harris et al. 2016; Grob et al. 2011). Such findings could be tentatively interpreted to imply that psilocybin facilitates more flexible, less constrained kinds of cognition (Carhart-Harris et al. 2016).

The 5-HT_2A_ receptors are widely distributed in in the brain and especially so in high-level prefrontal and associative cortex-regions important for learning and memory retrieval, this is likely to have important functional implications (Carhart-Harris and Nutt, 2017; Zhang and Stackman, 2015). For instance, postsynaptic 5-HT_2A_ receptor activation was shown to be associated with improvements in certain aspects of cognition (Gimpl et al., 1979; Harvey, 1996, 2003; Harvey et al., 2004, 2012; Romano et al., 2010; Zhang and Stackman, 2015; Zhang et al., 2016) as well as an extinction of previously learn response patterns (Zhang et al., 2013). However, it is important to note, that function of the 5-HT system remains ‘elusive’ given the inherent complexity of the serotonin system and more research has to be conducted in this regard to determine its function (Dayan and Huys, 2009; Carhart-Harris and Nutt, 2017).

While the assumption of a link between the use of psychedelics and an unconstrained brain state fits well with our findings on divergent thinking, it does not seem to be consistent with our observations on convergent thinking. Microdosing improved performance on the PCT, suggesting that it promotes convergent thinking. Note that this observation contrasts with previous findings by Kuypers and colleagues (2016), who reported that Ayahuasca, also a 5-HT_2A_ agonist, impaired performance on convergent thinking tasks. We believe this discrepancy could be a result of the difference in relative dosage. Kuypers and colleagues (2016) investigated participants after the intake of large doses of Ayahuasca, which is hardly comparable to the microdoses used in the present study. Previous research has shown a relationship between 5-HT_2A_ receptor activity and goal directed behavior likely due to indirect modulation of DA release (Vollenweider et al. 1999; Sakashita,et al. 2015; Dalley et al. 2002; Boureau and Dayan 2011). Dopamine-related adaptive behavior follows an inverted U-shape (van Velzen et al. 2014), suggesting that smaller doses, such as the microdoses ingested by the participants in our current study, are more likely to move participants towards the most efficient mid-zone of the performance function than higher doses do (see Akbari Chermahini and Hommel 2012, for an application of this rationale on the impact of dopaminergic manipulations on creativity). Indeed, based on self-reports, an online study by Fadiman and Krob (2017) suggests that microdosing could enhance motivation and focus, and reduce distractibility and procrastination—which seems consistent with our observation of improved convergent thinking.

These considerations suggest that microdoses truffles, and perhaps 5-HT_2A_ agonist in general, improve processes that are shared by convergent and divergent thinking—irrespective of the existing differences. Indeed, both convergent and divergent thinking tasks rely to some degree on persistence and top-down control and to some degree on unconstrained flexibility (Hommel, 2015). While convergent-thinking tasks emphasize persistence over flexibility, and divergent-thinking tasks emphasize flexibility over persistence, they both require participants to keep in mind particular search criteria, which they need to test against candidate items in memory (a skill that relies on persistence and top-down control), and to search through novel and often unfamiliar items considered for this test (a skill that relies on flexibility). The tasks thus present participants with a dilemma, which can only be solved by finding a reasonable balance between the antagonistic skills; that is, to be persistent and flexible at the same time, or at least in quick succession. Microdosing therefore might promote is the speed or smoothness of switching between persistence and flexibility—an ability that Mekern, Sjoerds, and Hommel (2018) refer to as “adaptivity”. Taken together, whereas large doses of psychedelics might induce an hyper-flexible mode of brain functioning, and possibly a breakdown of control (Carhart-Harris 2014), microdoses may be able to drive brain functioning towards an optimal, highly adaptive balance between persistence and flexibility.

### Limitations

It is important to consider the limitations of our study. The experiment was carried out “in the field”, which offers the benefits of studying more natural effects of microdosing, but also has the disadvantage of allowing less experimental control than in a laboratory setting. We used a quasi-experimental design, which limits some of the conclusions that can be drawn from our study. Neither the participants nor the researchers were blind to the manipulation and, as all participants consumed a microdose of psychedelic truffles, we were unable to collect data from a control group—which in this particular context would have been hard to conceive anyway. Relatedly, the absence of a placebo condition prevented any kind of randomization of the administered substance. Participants were not randomly selected due to the nature of the event, resulting in a self-selection bias, and limiting the generalizability of our findings. While we have argued that previous findings render the possibility of learning effects unlikely, it is also true that we were unable to quantify possible learning effects in the present study. Moreover, the absence of a control group did not allow us to identify and quantify possible placebo/expectation effects. Even though we have argued that this need not undermine our conclusions, future studies should definitely consider expectation/placebo effects, not only as a possible confound but also as a useful replacement of the actual drug. Another possible objection might be that mood might have contributed to our findings. Indeed, it is well-known that divergent thinking benefits from positive mood (Baas et al. 2008; Davis 2009), providing a possible confound regarding the effects we observed on our divergent thinking task. However, given that we also found positive effects for convergent thinking, which has not shown to benefit from positive mood, would not fit with a mood-related interpretation. Moreover, the well-established association between serotonin levels and depression (Neumeister 2003, Baldwin and Rudge 1995) suggests that mood should be considered the phenomenological expression of a particular serotonin level (arguably in combination with other factors), rather than an independent factor that can moderate the impact of changes in serotonin levels on cognitive performance. In retrospect, we also acknowledge that perceived strength of drug effect would have been a valuable measure to assess as it could have played some role in our outcome measures.

Furthermore, it has been shown that In contrast, however, positive mood negatively predicts convergent thinking performance, which in our study improved after ingestion of the psychedelic truffles. Future studies should seek to validate our findings using a lab-based randomized double-blind placebo controlled experimental designs and take the subjective strength of the experience into account as a covariate.

### Conclusion

Whereas large doses of psychedelics can introduce a range of undesirable side effects, microdoses of psychedelic substances might prove to be a promising alternative that could eliminate the risks of challenging experiences (sometimes referred to as a ''bad trips'') while maintaining the potential benefits of psychedelic substances on human emotion and cognition. The current naturalistic study is the first to quantitatively show that microdosing psychedelics could improve creative performance, possibly by means of inducing a state of unconstrained thought allowing for increased novel idea generation. We hope that our findings will stimulate further research into the beneficial effects of microdosing psychedelics. Apart from its benefits as a potential cognitive enhancement technique, microdosing could be further investigated for its therapeutic efficacy as to slow down cognitive decline or help individuals who suffer from rigid thought patterns or behavior such as individuals with depression or obsessive-compulsive disorder.

## Author Contributions

The study was set up together by all authors, who also contributed to data interpretation and writing the manuscript, and approved the final version of the manuscript. LP and DPL had the original idea, carried out the experiment, analyzed the data, and wrote a first draft of the article.

## Acknowledgements

We would like to thank the Psychedelic Society of the Netherlands for providing us with the opportunity to do the experiment at one of their events and Eliska Prochazkova for helping carrying out the experiment.

This research was partly funded by an Advanced Grant of the European Research Council (ERC-2015-AdG-694722) to BH.

## References

Aarts E, Roelofs A, Franke B, Rijpkema M, Fernández G, Helmich RC, Cools R (2010) Striatal dopamine mediates the interface between motivational and cognitive control in humans: Evidence from genetic imaging. Neuropsychopharmacology 35:1943–1951. doi: 10.1038/npp.2010.68

Akbari Chermahini S, Hickendorff M, Hommel B (2012) Development and validity of a Dutch version of the Remote Associates Task: An item-response theory approach. Think Ski Creat 7:177–186. doi: 10.1016/j.tsc.2012.02.003

Akbari Chermahini S, Hommel B (2010) The (b)link between creativity and dopamine: Spontaneous eye blink rates predict and dissociate divergent and convergent thinking. Cognition 115:458–465. doi: 10.1016/j.cognition.2010.03.007

Amabile TM (1996) Creativity in context: Update to the social psychology of creativity. Westview Press, Boulder, CO

Ashby FG, Isen AM, Turken AU (1999) A neuropsychological theory of positive affect and its influence on cognition. Psychol Rev 106:529–550. doi: 10.1037/0033-295X.106.3.529

Baas M, De Dreu CKW, Nijstad BA (2008) A Meta-Analysis of 25 Years of Mood-Creativity Research: Hedonic Tone, Activation, or Regulatory Focus? Psychol Bull 134:779–806. doi: 10.1037/a0012815

Baas M, Nevicka B, Ten Velden FS (2014) Specific mindfulness skills differentially predict creative performance. Personal Soc Psychol Bull 40:1092–1106. doi: 10.1177/0146167214535813

Baggot MJ (2015) Psychedelics and Creativity: A review of the quantitative literature. PeerJ Prepr 3:1–24

Baldwin D, Rudge S (1995). The role of serotonin in depression and anxiety. Int Clin Psychopharmacol 9:41–45

Bari A, Theobald DE, Caprioli D, Mar AC, Aidoo-Micah A, Dalley JW, Robbins TW (2010) Serotonin modulates sensitivity to reward and negative feedback in a probabilistic reversal learning task in rats. Neuropsychopharmacology 35:1290–1301. doi: 10.1038/npp.2009.233

Barrett FS, Griffiths RR (2017) Classic hallucinogens and mystical experiences: Phenomenology and neural vorrelates. In: Geyer MA, Ellenbroek BA, Marsden CA, Barnes TRE, L. Andersen S (eds) Current topics in behavioral neurosciences. Springer, Berlin, Heidelberg, pp 1–38

Barrett FS, Johnson MW, Griffiths RR (2017) Psilocybin in long-term meditators: Effects on default mode network functional connectivity and retrospective ratings of qualitative experience. Drug Alcohol Depend 171:e15–e16. doi: 10.1016/j.drugalcdep.2016.08.058

Bilker WB, Hansen JA, Brensinger CM, Richard J, Gur RE, Gur RC (2012) Development of abbreviated nine-item forms of the Raven’s Standard Progressive Matrices Test. Assessment 19:354–369. doi: 10.1177/1073191112446655

Bogenschutz MP, Forcehimes AA, Pommy JA, Wilcox CE, Barbosa P, Strassman RJ (2015) Psilocybin-assisted treatment for alcohol dependence: A proof-of-concept study. J Psychopharmacol 29:289–299. doi: 10.1177/0269881114565144

Boulougouris V, Glennon JC, Robbins TW (2008) Dissociable effects of selective 5-HT2A and 5-HT2C receptor antagonists on serial spatial reversal learning in rats. Neuropsychopharmacology 33:2007–2019. doi: 10.1038/sj.npp.1301584

Boureau YL, Dayan P (2011) Opponency revisited: Competition and cooperation between dopamine and serotonin. Neuropsychopharmacology 36:74–97

Callaway JC, Grob CS (1998) Ayahuasca preparations and serotonin reuptake inhibitors: A potential combination for severe adverse interactions. J Psychoactive Drugs 30:367–369. doi: 10.1080/02791072.1998.10399712

Carhart-Harris RL, Bolstridge M, Rucker J, Day CMJ, Erritzoe D, Kaelen M, Bloomfield M, Rickard JA, Forbes B, Feilding A, Taylor D, Pilling S, Curran VH, Nutt DJ (2016) Psilocybin with psychological support for treatment-resistant depression: An open-label feasibility study. The Lancet Psychiatry 3:619–627. doi: 10.1016/S2215-0366(16)30065-7

Carhart-Harris RL, Kaelen M, Bolstridge M, Williams TM, Williams LT, Underwood R, … and Nutt DJ (2016). The paradoxical psychological effects of lysergic acid diethylamide (LSD). Psychol Med 46:1379–1390

Carhart-Harris RL, Leech R, Hellyer PJ, Shanahan M, Feilding A, Tagliazucchi E, Chialvo DR, Nutt D, Axmacher N, Brooks SJ, Maclean K, Hopkins J (2014) The entropic brain: A theory of conscious states informed by neuroimaging research with psychedelic drugs. Front Hum Neurosci 8:20. doi: 10.3389/fnhum.2014.00020

Carhart-Harris RL, and Nutt DJ (2017). Serotonin and brain function: a tale of two receptors. J Psychopharmacol 31: 1091–1120.

Catlow BJ, Jalloh A, Sanchez-Ramos J (2016) Hippocampal neurogenesis: Effects of psychedelic drugs. In: Neuropathology of drug addictions and substance misuse. Elsevier, pp 821–831

Clarke HF, Dalley JW, Crofts HS, Robbins TW, Roberts AC (2004) Cognitive inflexibility after prefrontal serotonin depletion. Science (80-) 304:878–880. doi: 10.1126/science.1094987

Clarke HF, Walker SC, Dalley JW, Robbins TW, Roberts AC (2007) Cognitive inflexibility after prefrontal serotonin depletion is behaviorally and neurochemically specific. Cereb Cortex 17:18–27. doi: 10.1093/cercor/bhj120

Colzato LS, de Haan AM, Hommel B (2015) Food for creativity: Tyrosine promotes deep thinking. Psychol Res 79:709–714. doi: 10.1007/s00426-014-0610-4

Cooke R (2017) How dropping acid saved my life. The Guardian. https://www.theguardian.com/global/2017/jan/08/how-dropping-acid-saved-my-life-ayelet-waldman-books-depression. Accessed 25 Dec 2017

Cools R, D’Esposito M (2009) Dopaminergic modulation of flexible control in humans. In: Bjorklund A, Dunnett SB, Iversen LL, Iversen SD (eds) Dopamine handbook. Oxford University Press, Oxford

Cools R, D’Esposito M (2011) Inverted-U-shaped dopamine actions on human working memory and cognitive control. Biol. Psychiatry 69:e113–e125

Dalley JW, Theobald DE, Eagle DM, Passetti F, Robbins TW (2002) Deficits in impulse control associated with tonically-elevated serotonergic function in rat prefrontal cortex. Neuropsychopharmacology 26:716–728. doi: 10.1016/S0893-133X(01)00412-2

Darvas M, Palmiter RD (2011) Contributions of striatal dopamine signaling to the modulation of cognitive flexibility. Biol Psychiatry 69:704–707. doi: 10.1016/j.biopsych.2010.09.033

Davis MA (2009) Understanding the relationship between mood and creativity: A meta-analysis. Organ Behav Hum Decis Process 108:25–38. doi: 10.1016/j.obhdp.2008.04.001

De Dreu CKW, Baas M, Nijstad BA (2008) Hedonic tone and activation level in the mood-creativity link: Toward a dual pathway to creativity model. J Pers Soc Psychol 94:739–756. doi: 10.1037/0022-3514.94.5.739

dos Santos RG, Osório FL, Crippa JAS, Riba J, Zuardi AW, Hallak JEC (2016) Antidepressive, anxiolytic, and antiaddictive effects of ayahuasca, psilocybin and lysergic acid diethylamide (LSD): A systematic review of clinical trials published in the last 25 years. Ther Adv Psychopharmacol 6:193–213. doi: 10.1177/2045125316638008

Dayan P and Huys QJ (2009) Serotonin in affective control. Annu Rev Neurosci 32:95–126

Dreisbach G, Goschke T (2004) How positive affect modulates cognitive control: Reduced perseveration at the cost of increased distractibility. J Exp Psychol Learn Mem Cogn 30:343–353. doi: 10.1037/0278-7393.30.2.343

Erowid (2017) Psilocybin mushrooms effects. The Vaults of Erowid. https://erowid.org/plants/mushrooms/mushrooms_effects.shtml. Accessed 25 Dec 2017

Fadiman J (2011) The Psychedelic Explorer’s Guide: Safe, Therapeutic, and Sacred Journeys. Inner Traditions Bear and Company. Rochester, NY.

Fadiman J, Krob S (2017) Microdosing: The phenomenon, research results, and startling surprises. Lecture presented at the Psychedelic Science 2017 conference, Oakland, CA

Friston K, Kilner J, Harrison L (2006) A free energy principle for the brain. J Physiol Paris 100:70–87. doi: 10.1016/j.jphysparis.2006.10.001

Farah MJ, Haimm C, Sankoorikal G, and Chatterjee A (2009) When we enhance cognition with Adderall, do we sacrifice creativity? A preliminary study. Psychopharmacol, 202: 541–547.

Geyer MA, Vollenweider FX (2008) Serotonin research: Contributions to understanding psychoses. Trends Pharmacol. Sci. 29:445–453

Gimpl MP, Gormezano I and Harvey JA (1979) Effects of LSD on learning as measured by classical conditioning of the rabbit nictitating membrane response. J Pharmacol Exp Ther 208: 330–334.

Glatter R (2015) LSD microdosing: The new job enhancer in Silicon Valley. Forbes. https://www.forbes.com/sites/robertglatter/2015/11/27/lsd-microdosing-the-new-job-enhancer-in-silicon-valley-and-beyond/#109a749b188a. Accessed 25 Dec 2017

Gregorie C (2016) Everything you wanted to know about microdosing (but were afraid to ask). The Huffington Post. http://www.huffingtonpost.com/entry/psychedelic-microdosing-research_us_569525afe4b09dbb4bac9db8. Accessed 25 Dec 2017

Griffiths RR, Richards WA, McCann U, Jesse R (2006) Psilocybin can occasion mystical-type experiences having substantial and sustained personal meaning and spiritual significance. Psychopharmacology 187:268–283. doi: 10.1007/s00213-006-0457-5

Grob CS, Danforth AL, Chopra GS, Hagerty M, McKay CR, Halberstad AL, Greer GR (2011) Pilot study of psilocybin treatment for anxiety in patients with advanced-stage cancer. Arch Gen Psychiatry 68:71–78. doi: 10.1001/archgenpsychiatry.2010.116

Gartz J, Allen JW, Merlin MD (1994) Ethnomycology, biochemistry, and cultivation of Psilocybe samuiensis Guzman, Bandala and Allen, a new psychoactive fungus from Koh Samui, Thailand. J Ethnopharmacol, 43:73–80

Guilford JP (1967) The nature of human intelligence. McGraw-Hill, NY

Harvey JA, Quinn JL, Liu R, et al. (2004) Selective remodeling of rabbit frontal cortex: relationship between 5-HT2A receptor density and associative learning. Psychopharmacology (Berl) 172: 435–442.

Harvey JA, Schlosberg AJ and Yunger LM (1975) Behavioral correlates of serotonin depletion. Fed Proc 34: 1796–1801.

Harvey ML, Swallows CL and Cooper MA (2012) A double dissociation in the effects of 5-HT2A and 5- HT2C receptors on the acquisition and expression of conditioned defeat in Syrian hamsters. Behav Neurosci 126: 530–537.

Hajkova K, Jurasek B, Sykora D, Palenicek T, Miksatkova P, Kuchar M (2016) Salting-out-assisted liquid-liquid extraction as a suitable approach for determination of methoxetamine in large sets of tissue samples. Anal Bioanal Chem 408:1171–1181. doi: 10.1007/s00216-015-9221-1

Halberstadt AL (2015) Recent advances in the neuropsychopharmacology of serotonergic hallucinogens. Behav. Brain Res. 277: 99–120.

Hasler F, Grimberg U, Benz MA, Huber T, and Vollenweider, FX (2004). Acute psychological and physiological effects of psilocybin in healthy humans: a double-blind, placebo-controlled dose-effect study. Psychopharmacol 172: 145–156.

Haluk DM, Floresco SB (2009) Ventral striatal dopamine modulation of different forms of behavioral flexibility. Neuropsychopharmacology 34:2041–2052. doi: 10.1038/npp.2009.21

Harman WW, Fadiman J (1970) Selective enhancement of specific capacities through psychedelic training. In: Aaronson B, Osmond H (eds) Psychedelics: The uses and implications of psychedelic drugs. Anchor Books, Garden City, NY, pp 239–256

Harman WW, McKim RH, Mogar RE, Fadiman J, Stolaroff MJ (1966) Psychedelic agents in creative problem-solving: A pilot study. Psychol Rep 19:211–227. doi: 10.2466/pr0.1966.19.1.211

Harvey JA (2003) Role of the serotonin 5-HT2A receptor in learning. Learn Mem 10:355–362. doi: 10.1101/lm.60803

Harvey JA (1995) Serotonergic regulation of associative learning. Behav Brain Res 73:47–50. doi: 10.1016/0166-4328(96)00068-X

Hasler F, Grimberg U, Benz MA, Huber T, Vollenweider FX (2004) Acute psychological and physiological effects of psilocybin in healthy humans: a double-blind, placebo-controlled dose-effect study. Psychopharmacology 172: 145–156.

Hollister LE (1968) Chemical psychoses: LSD and related drugs. Charles C. Thomas, Springfield, IL

Hommel B (2012) Convergent and divergent operations in cognitive search. In: Todd PM, Hills TT, Robbins TW (eds) Cognitive search: Evolution, algorithms, and the brain, vol. 9. MIT Press, Cambridge, MA, pp 215–230

Hommel B (2015) Between persistence and flexibility: The Yin and Yang of action control. In: Elliot AJ (ed) Advances in motivation science, vol. 2. Elsevier, NY, pp 33–67

Hommel B, Colzato LS (2017) The social transmission of metacontrol policies: Mechanisms underlying the interpersonal transfer of persistence and flexibility. Neurosci. Biobehav. Rev. 81:43–58

Hurks PPM, Hendriksen J, Dek JE, Kooij AP (2010) De nieuwe Wechsler kleuterintelligentietest voor 2:6-7:11 jarigen. Tijdschrift voor Neuropsychologie 2:40–51

Jensen A (1969) How much can we boost IQ and scholastic achievement. Harv Educ Rev 39:1–123. doi: 10.17763/haer.39.1.l3u15956627424k7

Johnson MW, Garcia-Romeu A, Cosimano MP, Griffiths RR (2014) Pilot study of the 5-HT2AR agonist psilocybin in the treatment of tobacco addiction. J Psychopharmacol 28:983–992. doi: 10.1177/0269881114548296

Kowal MA, Hazekamp A, Colzato LS, van Steenbergen H, van der Wee NJ, Durieux J., … and Hommel B (2015). Cannabis and creativity: Highly potent cannabis impairs divergent thinking in regular cannabis users. Psychopharmacol, 232: 1123–1134.

Krebs TS, Johansen Pør (2012) Lysergic acid diethylamide (LSD) for alcoholism: Meta-analysis of randomized controlled trials. J. Psychopharmacol. 26:994–1002

Kuypers KPC, Riba J, de la Fuente Revenga M, Barker S, Theunissen EL, Ramaekers JG (2016) Ayahuasca enhances creative divergent thinking while decreasing conventional convergent thinking. Psychopharmacology (Berl) 233:3395–3403. doi: 10.1007/s00213-016-4377-8

Leonard A (2015) How LSD microdosing became the hot new business trip. Rolling Stone. http://www.rollingstone.com/culture/features/how-lsd-microdosing-became-the-hot-new-business-trip-20151120. Accessed 25 Dec 2017

Lippelt DP, Hommel B, Colzato LS (2014) Focused attention, open monitoring and loving kindness meditation: Effects on attention, conflict monitoring, and creativity - A review. Front. Psychol. 5:1083

MacLean KA, Johnson MW, Griffiths RR (2011) Mystical experiences occasioned by the hallucinogen psilocybin lead to increases in the personality domain of openness. J Psychopharmacol 25:1453–1461. doi: 10.1177/0269881111420188

Matias S, Lottem E, Dugué GP, Mainen ZF (2017) Activity patterns of serotonin neurons underlying cognitive flexibility. Elife 6:1–24. doi: 10.7554/eLife.20552

Mednick S (1962) The associative basis of the creative process. Psychol Rev 69:220–232. doi: 10.1037/h0048850

Moreno FA, Wiegand CB, Taitano EK, Delgado PL (2006) Safety, tolerability, and efficacy of psilocybin in 9 patients with obsessive-compulsive disorder. J Clin Psychiatry 67:1735–40

Neumeister A (2003). Tryptophan depletion, serotonin, and depression: Where do we stand?. Psychopharmacol Bull, 37:99–115

Oberhaus D (2017) 400 people microdosed LSD for a month in the name of science. Vice Magazine. https://motherboard.vice.com/en_us/article/mgy573/400-people-microdosed-lsd-for-a-month-in-the-name-of-science. Accessed 25 Dec 2017

Quitkin F, Rifkin A, Klein DF (1979) Monoamine oxidase inhibitors. A review of antidepressant effectiveness. Arch Gen Psychiatry 36:749–60

Raven J (1938) Raven’s progressive matrices (1938): Sets A, B, C, D. Australian Council for Educational Research, Melbourne

Raven J (2000) The Raven’s Progressive Matrices: Change and stability over culture and time. Cogn Psychol 41:1–48. doi: 10.1006/cogp.1999.0735

Reuter M, Roth S, Holve K, Hennig J. Identification of the first candidate genes for creativity: A pilot study. Brain Res. 2006; 1069:190–7. pmid: 16403463

Riba J, Romero S, Grasa E, Mena E, Carrió I, Barbanoj MJ (2006) Increased frontal and paralimbic activation following ayahuasca, the pan-amazonian inebriant. Psychopharmacology (Berl) 186:93–98. doi: 10.1007/s00213-006-0358-7

Ritter SM, Mostert N (2016) Enhancement of creative thinking skills using a cognitive-based creativity training. J Cogn Enhanc 1:243–253. doi: 10.1007/s41465-016-0002-3

Romano AG, Quinn JL, Li L, et al. (2010) Intrahippocampal LSD accelerates learning and desensitizes the 5- HT(2A) receptor in the rabbit. Psychopharmacology (Berl) 212:441–448

Sahakian B (2017) LSD microdosing is trending in Silicon Valley but can actually make you more creative. IFLScience. http://www.iflscience.com/health-and-medicine/lsd-microdosing-is-trendingin-silicon-valley-but-can-actually-make-you-more-creative/. Accessed 25 Dec 2017

Sakashita Y, Abe K, Katagiri N, Kambe T, Saitoh T, Utsunomiya I, Horiguchi Y, Taguchi K (2015) Effect of psilocin on extracellular dopamine and serotonin levels in the mesoaccumbens and mesocortical pathway in awake rats. Biol Pharm Bull 38:134–138. doi: 10.1248/bpb.b14-00315

Schafer G, Feilding A, Morgan CJ, Agathangelou M, Freeman TP and Curran, HV (2012). Investigating the interaction between schizotypy, divergent thinking and cannabis use. Consc and Cogn, 21: 292–298.

Schwarz N (1999) Self-reports: How the questions shape the answers. Am Psychol 54:93–105. doi: 10.1037/0003-066X.54.2.93

Schwarz K, Pfister R, Büchel C. Rethinking Explicit Expectations: Connecting Placebos, Social Cognition, and Contextual Perception. Trends in Cognitive Sciences. 2016.

Senior J (2017) Ayelet Waldman’s better living through LSD. The New York Times. https://www.nytimes.com/2017/01/11/books/review-a-really-good-day-ayelet-waldmans-better-living-through-lsd.html. Accessed 25 Dec 2017

Sessa B (2008) Is it time to revisit the role of psychedelic drugs in enhancing human creativity? J. Psychopharmacol. 22:821–827

Solon O (2016) Would you take LSD to give you a boost at work? WIRED takes a trip inside the world of microdosing. WIRED. http://www.wired.co.uk/article/lsd-microdosing-drugs-silicon-valley. Accessed 25 Dec 2017

Sokolov EN (1960) Neuronal models and the orienting reflex. In: Brazier MAB (ed) The central nervous system and behavior. JosiahMacy, Jr. Foundation, NY, pp 187–276

Sternberg RJ (2008) Increasing fluid intelligence is possible after all. Proc Natl Acad Sci 105:6791–6792. doi: 10.1073/pnas.0803396105

Sternberg RJ, Lubart TI (1999) The concept of creativity: Prospects and paradigms. In: Sternberg RJ (ed) Handbook of creativity. Cambridge University Press, London, pp 3–16

Stevenson Claire E., Kleibeuker Sietske W., de Dreu Carsten K. W., Crone Eveline A. (2014). Training creative cognition: adolescence as a flexible period for improving creativity. Frontiers in Human Neuroscience, 8: 827.

Tolman EC, Brunswik E (1935) The organism and the causal texture of the environment. Psychol Rev 42:43–77. doi: 10.1037/h0062156

Tylš F, Páleníček T, Horáček J (2014) Psilocybin - Summary of knowledge and new perspectives. Eur. Neuropsychopharmacol. 24:342–356

van Velzen LS, Vriend C, de Wit SJ, van den Heuvel OA (2014) Response inhibition and interference control in obsessive-compulsive spectrum disorders. Front Hum Neurosci 8:419. doi: 10.3389/fnhum.2014.00419

Vollenweider FX, Geyer MA (2001) A systems model of altered consciousness: Integrating natural and drug-induced psychoses. Brain Res Bull 56:495–507

Vollenweider FX, Kometer M (2010) The neurobiology of psychedelic drugs: Implications for the treatment of mood disorders. Nat Rev Neurosci 11:642–651. doi: 10.1038/nrn2884

Vollenweider F, Vontobel P, Hell D, Leenders KL (1999) 5-HT modulation of dopamine release in basal ganglia in psilocybin-induced psychosis in man—A PET study with [11C]raclopride. Neuropsychopharmacology 20:424–433. doi: 10.1016/S0893-133X(98)00108-0

Vosburg SK (1998) The effects of positive and negative mood on divergent-thinking performance. Creat Res J 11:165–172. doi: 10.1207/s15326934crj1102_6

Wallas G (1926) The art of thought. Harcourt Brace, NY

Wechsler D (2003) The Wechsler intelligence scale for children, 4th edn. Pearson, London

Zabelina DL, Robinson MD (2010) Creativity as flexible cognitive control. Psychol Aesthetics, Creat Arts 4:136–143. doi: 10.1037/a0017379

Zabelina, D.L., Colzato, L.S., Beeman, M., Hommel, B. (2016). Dopamine and the creative mind: Individual differences in creativity are predicted by interactions between dopamine genes DAT and COMT. PLoS One. 11: e0146768.

Zegans LS, Pollard JC, Brown D (1967) The effects of LSD-25 on creativity and tolerance to regression. Arch Gen Psychiatry 16:740–749. doi: 10.1001/archpsyc.1967.01730240096014

Zhang G, Asgeirsdottir HN, Cohen SJ, et al. (2013) Stimulation of serotonin 2A receptors facilitates consolidation and extinction of fear memory in C57BL/6J mice. Neuropharmacology 64: 403–413.

Zhang G, Cinalli D, Cohen SJ, et al. (2016) Examination of the hippocampal contribution to serotonin 5- HT2A receptor-mediated facilitation of object memory in C57BL/6J mice. Neuropharmacology 109: 332–340.

Zhang G and Stackman RW (2015) The role of serotonin 5-HT2A receptors in memory and cognition. Front Pharmacol 6:225.

Zhang ZW (2003) Serotonin induces tonic firing in layer V pyramidal neurons of rat prefrontal cortex during postnatal development. J Neurosci 23: 3373–3384.

